# Differential co-expression network analysis with DCoNA reveals isomiR targeting aberrations in prostate cancer

**DOI:** 10.1101/2022.06.28.497906

**Authors:** Anton Zhiyanov, Narek Engibaryan, Stepan Nersisya, Maxim Shkurnikov, Alexander Tonevitsky

## Abstract

We developed DCoNA – a statistical tool that allows one to identify pair interactions, which correlation significantly changes between two conditions. Comparing DCoNA with the state-of-the-art analog, we showed that DCoNA is a faster, more accurate, and less memory-consuming tool. We applied DCoNA to prostate mRNA/miRNA-seq data collected from The Cancer Genome Atlas (TCGA) and compared predicted regulatory interactions of miRNA isoforms (isomiRs) and their target mRNAs between normal and cancer samples. As a result, almost all highly expressed isomiRs lost negative correlation with their targets in prostate cancer samples compared to ones without the pathology. One exception to this trend was the canonical isomiR of hsa-miR-93-5p acquiring cancer-specific targets. Further analysis showed that cancer aggresiveness increased with the expression of this isomiR in both TCGA primary tumor samples and 153 blood plasma samples of own patients’ cohort analyzed by miRNA microarrays.

## Introduction

MicroRNAs (miRNAs) are small non-coding RNA molecules that regulate gene expression by binding to mRNA targets and initiating their degradation or inhibiting their translation [1]. It was shown that miRNAs could act as tumor suppressors and oncogenes in human cancers. For instance, the members of the miR-200 family (miR-200a, miR-200b, miR-200c, miR-141, and miR-449) inhibit metastasis of various cancers by targeting ZEB1 and ZEB2, the master regulators of epithelial-to-mesenchymal transition [2, 3].

After transcription, a pri-miRNA hairpin is processed by Drosha in the cell nucleus. The resulting pre-miRNA molecule is then moved to the cytoplasm, where the miRNA duplex is formed after Dicer cleavage. One strand of this duplex is preferentially incorporated into RISC complex, becoming the mature microRNA [4]. Heterogeneous enzymatic cleavage by Drosha and Dicer leads to the length variation at 5’- and 3’-ends of miRNAs, which results in the generation of miRNA isoforms (isomiRs) (Figure 1 A) [5]. The importance of isomiRs can be explained by the fact that different isomiRs of the same miRNA may have different target genes if the seed region of the molecule (nucleotides 2-7 counting from the 5’-end) is affected (Figure 1 B). For example, we recently showed that undesired 5’-isomiR of ELOVL5 shRNA (which can be considered as an exogenous miRNA) induces significant miRNA-like off-target effects after transduction in MDA-MD-231 cells [6].

**Figure 1:**
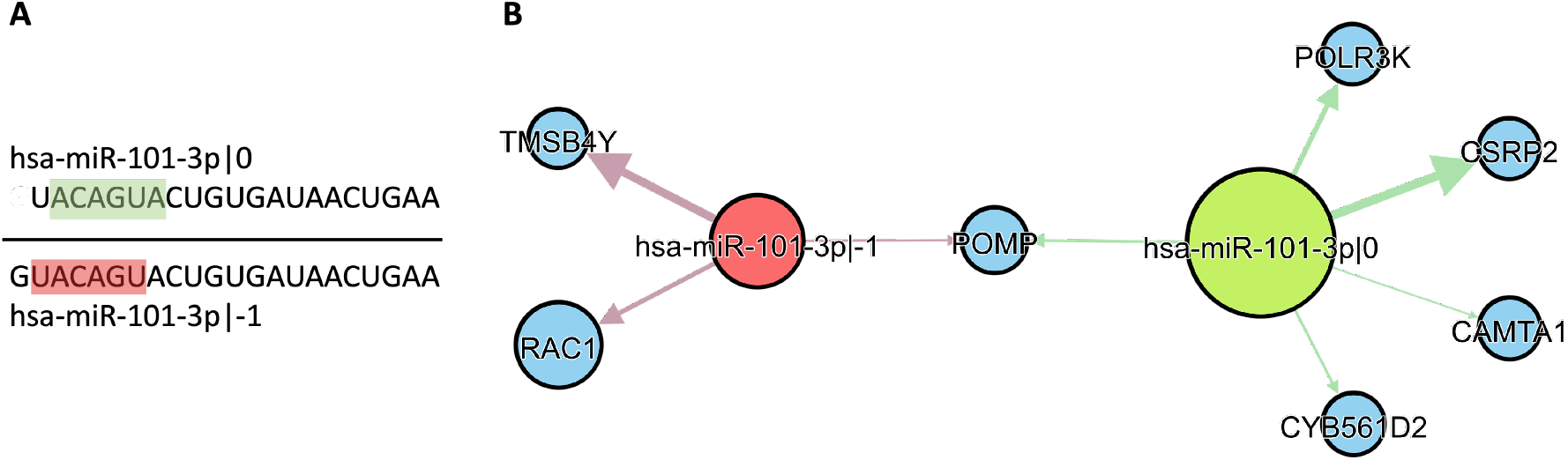
**A**: Two isomiRs of hsa-miR-101-3p have different seed regions (2-7 nts) highlighted red and green. **B**: These isomiRs potentionally regulate different genes in prostate cancer. The width of edges represents an absolute value of Spearman correlation between an isomiR and its target expression levels. All correlations are less than —0.3.

High-throughput RNA sequencing is widely used to study the biological role of miRNAs and isomiRs in the process of interest. The conventional differential expression analysis highlights those isomiRs and their target mRNAs, which average expression levels significantly differ between two conditions [7, 8]. Such molecules can be further considered as indicators of pathological processes or used to build prognostic models. However, differential expression methods do not detect changes in molecular regulation in case if average expression levels of regulatory and target molecules are not changed (Figure 2). Such a situation was described, e.g., by Keller et al [9]. The authors showed that p16 and the group of cyclins were negatively co-expressed in lean mice but positively co-expressed in obese mice, suggesting an obesity-related regulation of the cell cycle pathway. Notably, p16 and many of the cyclins were not differentially expressed between the lean and obese mice and would have therefore been missed by differential expression analysis.

**Figure 2:**
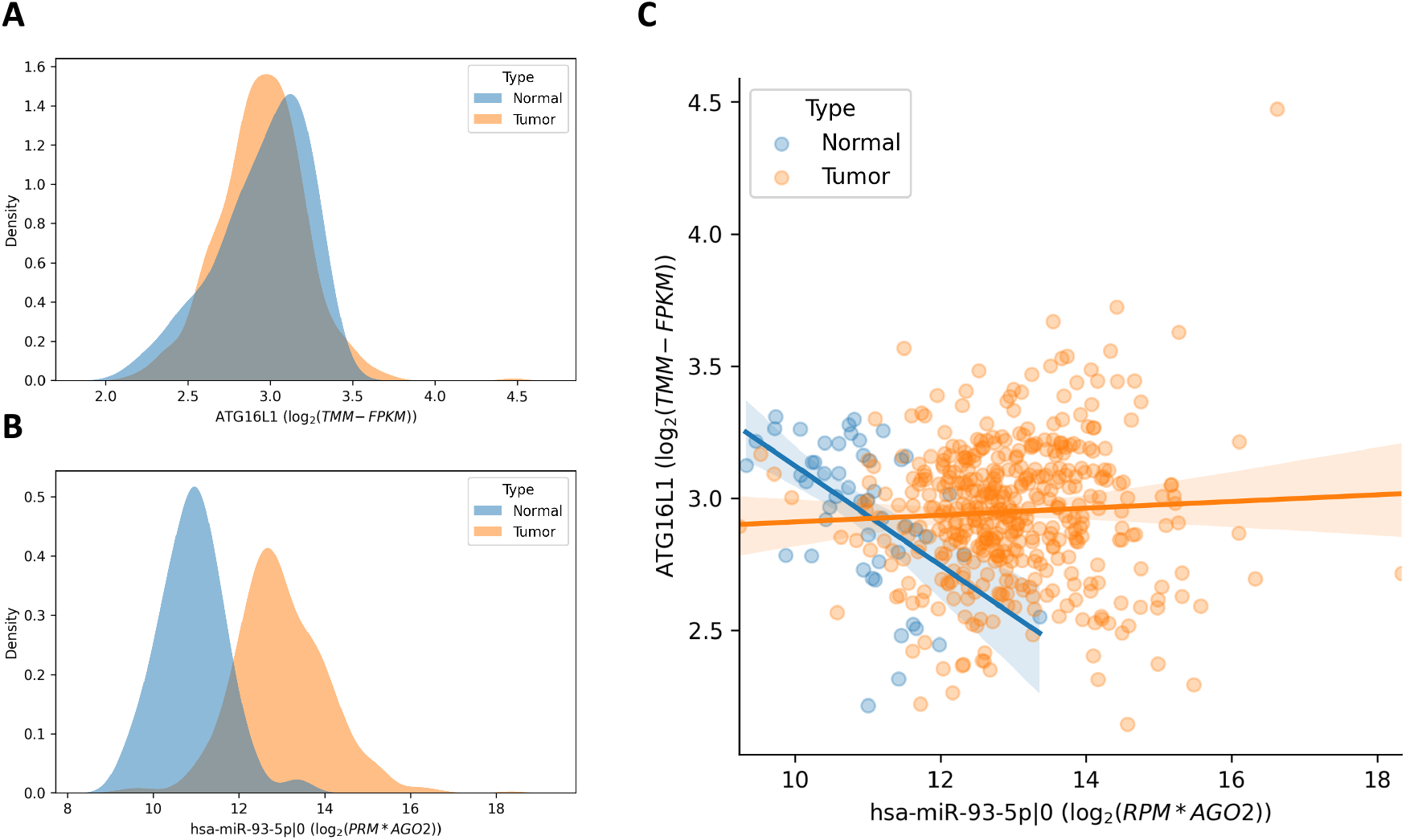
**A**: Expression of ATG16L1 gene does not significantly change in prostate cancer compared with a normal state, while expression of hsa-miR-93-5p|0 changes (**B**). **C**: Moreover, the mutual expression of the gene and its regulatory isomiR changes: Spearman correlation of the molecules’ expression increases from —0.53 to 0.04.

So, we see that, in contrast to differential expression analysis, differential co-expression analysis aims to detect pairs or clusters which mutual expression changes between two conditions. For example, such changes may indicate a loss of regulation between miRNA and its mRNA target due to mutations in the binding region. In general, differential co-expression analysis can be divided into two major classes. The first class deals with pairwise co-expression and detects pairs whose interaction significantly changed between conditions. Several metrics like Pearson or Spearman correlation and Mutual information are used to evaluate the co-expression level numerically. Comparing values of these metrics under the conditions, one can conclude that the co-expression of a particular pair significantly changed. As an example of tools computing correlation to detect co-expression alterations we mention DGCA [10] and DiffCorr [11]. Here, the authors used correlation-based Fisher *z*-statistic [12] and tested a hypothesis that correlations under the conditions are equal. The studies [11, 13] categorized pairs into all possible paired correlation scenarios, that among other things, allowed us to identify pairs that experience no correlation in one condition but become correlated in another. As an example of non-correlation methods, we mention PMINR [14], where authors used mutual information and build a regression model to detect pairwise interactions related to a disease.

The second type of analysis detects co-expressed gene modules or clusters based on the similarity of their gene expression in each condition. Examples of such studies are WGCNA [15] and MEGENA [16], computing module overlap statistics between conditions [17] or the average modular differential connectivity [18].

In this paper, we study the differences in co-expression between 5’-isomiRs and their mRNA targets in normal prostate (“Normal”) and prostate cancer (“Tumor”). For that, we decided to develop our tool (DCoNA). It was based on the same principles as DGCA, with some changes. Like DGCA, DCoNA computes Fisher *z*-statistic to check the correlation equality hypothesis. However, DCoNA differs from the previous works in the following directions:

### Network analysis

While omics databases contain expression of thousands of mRNAs, microRNAs, and other molecules, only a few molecule interactions among all pairs are biologically meaningful. So, DCoNA can be applied not only to the complete network consisting of all pairs of molecules but also to a predefined set of interactions. On the one hand, this approach is more reasonable than the complete network analysis since multiple hypothesis adjustment reduces the significance of the conclusion regarding a particular interaction. On the other hand, it does not waste computational time on uniteresting pairs.

### Both analytic and permutation hypothesis testing approaches

As discussed in the next section, we use a statistical test to compute p-values under the correlation equality hypothesis analytically for both Pearson and Spearman correlation coefficients. Despite the theoretical guarantees, the analytical approach may be inaccurate if the sample sizes of two groups (“Normal” and “Tumor”) are small. DCoNA uses the permutation method to tackle this problem: it shuffles groups’ labels, recomputes the statistics, and compares them with initial ones.

### Aggregation scores

Note that pairwise correlation analysis may lack interpretability: some mediate correlation alterations related to a particular molecule may be discarded due to the multiple hypothesis adjustment. However, these changes may highlight the molecule as one with many reduced negative correlations. So, DCoNA computes scores aggregating *z*-statistics of targets and checks their significance using the permutation method.

### Efficiency

DCoNA was implemented using C++ language, while its analogs are based on R, Python, and other interpreted programming languages. So, DCoNA works significantly faster than the previous tools. Moreover, DCoNA supports an efficient parallel mode that further reduces computational time without using any additional memory.

### Application

We executed DCoNA on TCGA-PRAD dataset using predicted 5’-isomiR targets. We also illustrated DCoNA characteristics on synthetic datasets and compared them with state-of-the-art DGCA.

## Materials and methods

### Statistical framework

#### Correlation equivalence test

Let us assume that we have two independent paired samples 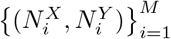 and 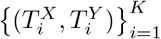 taken under two conditions: the first sample can be considered as expression levels of two genes “X” and “Y” taken from “Normal” group, and the second one consists of the expressions of the same genes taken from “Tumor” group. The “Tumor” and “Normal” notation is used only for the simplicity and the method could be applied to any two conditions. Denote by *P*(*N*),*P*(*T*) Pearson correlation coefficients of “X” and “Y” genes computed in “Normal” and “Tumor” groups, respectively. Similarly, we denote corresponding Spearman correlation coefficients by *S*(*N*) and *S*(*T*).

Assuming that pairs 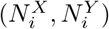 are bivariate normally distributed (BVN) with 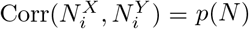, Fisher transformation *z*(*P*(*N*)) = arctanh(*P*(*N*)) of the correlation coefficient *P*(*N*) is asymptotically normal with mean *z*(*p*(*N*)) and asymptotic variance independent (as function) of *p*(*N*) [19]. Moreover, for non-BVN pairs, the asymptotic normality still holds, but the asymptotic variance may depend on *p*(*N*) [19]. Based on this idea, several methods were proposed to estimate the distribution of *z*(*P*(*N*)) and *z*(*S*(*N*)) for a fixed sample size. Following the works [20, 21], the transformed correlation coefficients can be approximated by the normal distributions with the means *z*(*p*(*N*)), *z*(*s*(*N*)) and the asymptotic variances

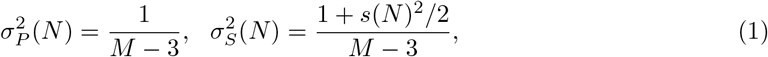

for Pearson and Spearman correlations, respectively.

In the co-expression difference analysis we test a hypothesis that correlations between genes “X” and “Y” are the same in “Normal” and “Tumor” groups, i.e. *p*(*N*) = *p*(*T*). Under the hypothesis, the statistic

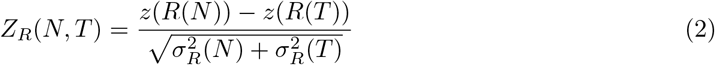

is asymtotically normal with zero mean and unit variance, where *R* is Pearson (*P*) or Spearman (*S*) correlation coefficient, and 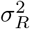 is the corresponding asymptotic variance. In the case of Pearson correlation the mentioned statistic does not depend on the correlation *p*(*N*) that allows us to compute the test *p*-value *P*(*ξ* > *z_P_*(*N,T*)), where *z_P_*(*N,T*) is the value of the statistic *Z_P_*(*N,T*) computed by the observed expressions and is a standard normal random variable. On the contrary, as follows from equations (1) and (2), the test statistic *Z_S_*(*N,T*) depends on the correlation *s*(*N*). Thus we use the value

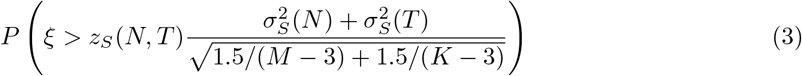

to upper bound the test p-value *P* (*ξ* > *z_S_*(*N, T*)).

#### Bootstrap p-value

However, the discussed analytical approach has several limitations. In general, we do not know in advance whether the distribution of the samples is BVN. Moreover, the asymptotic normality does not define the sample size starting from which the asymptotic approximation can be used. Thus we use the *permutation test* to compute p-value: we shuffle groups’ labels (“Normal”/”Tumor”), preserving the size of each group; recompute the value of the statistic *z_R_*(*N,T*); find a proportion of the values that exceed the value computed on the original group labeling. This procedure is correct under the assumption that the label permutations allow us to sample the value of the statistic *Z_R_*(*N,T*) under the hypothesis (*p*(*N*) = *p*(*T*) or *s*(*N*) = *s*(*T*), but maybe unknown in advance).

In practice, we simultaneously test the hypothesis on correlation equivalence among several pairs of interactions (“X” and “Y”). Thus, we adjust the obtained p-values using Benjamini-Hocheberg [22] correction method. We note that this correction defines a minimum number of permutations: if we simultaneously test *L* hypotheses, we should make at least *L/α* permutations to get the adjusted p-value with precision capable of rejecting the hypothesis with significance level *α*. DCoNA uses this minimum number as a default number of permutations.

### How to aggregate correlation differences

We note that the pairwise correlation analysis may lack interpretability: if the number of tested pairs is large, it may be difficult to extract source molecules (the first elements of the pairs) with a large percent of changed correlations. So, it may be helpful to group pairs with changed correlations by the source molecules and find overrepresented groups using the hypergeometric test, i.e., check that the number of significantly (the adjusted *p*-value < 0.05) changed correlations *s_G_* associated with a particular source molecule *G* is uniformly distributed among all significant changes. More precisely, DCoNA assumes that *s_G_* has hypergeometric distribution *H*(*N, S,n_G_*), where *n_G_* is the number of interactions with the molecule *G* in the predefined interaction network, *N* = ∑_*g*_ *n_g_* is the total number of interactions in the network and *S* = ∑_*g*_ *s_g_* is the total number of significantly changed correlations.

However, due to the multiple hypothesis adjustment, the hypergeometric test may lose some mediate correlation alterations associated with a particular source molecule. Moreover, finding directions of alterations associated with the molecule can be interesting. So, DCoNA can also compute mean, median and other quantiles of *z_R_*(*N,T*) statistics associated with the molecule and test the significance of this aggregated score using the label permutation test: it computes p-value as the fraction of permutations such that the absolute value of the score after permutation is greater than the value calculated on the initial labeling. Using this procedure, DCoNA tests a hypothesis that the aggregated score equals zero, i.e., the molecule has the same correlations under both conditions.

## Usage modes

DCoNA’s pipeline is shown in Figure 8 of “Supplementary materials”. As we have already discussed, DCoNA was designed to test the hypothesis on correlation equivalence for a predefined list of source and target pairs (“network.ztest” mode). However, DCoNA can also be used in the complete-network regime when the list is not given (“exhaustive.ztest” mode). In this regime, DCoNA tests the hypothesis for all possible pairs of molecules from expression data.

Aside from the hypothesis testing, DCoNA can be used to test that significantly altered correlations of a particular source molecule are overrepresented among all significantly changed correlations (“network(exhaustive).hypergeom” mode). Also, DCoNA can compute mean, median, and other quantiles of *z*-statistics associated with a particular molecule and its targets to determine a trend in correlation changes (“network(exhaustive).zscore”). All necessary parameters can be defined in a configuration file.

A more detailed description of DCoNA usage modes, the configuration file structure, and data format can be found at https://github.com/zhiyanov/DCoNA.

## Implementation

Pairwise differential co-expression analysis is computationally challenging. Indeed, let us assume that we have expression data of shape n × *m,* where n is the number of genes, and *m* is the sample size. Suppose we test all gene-gene pairs for equality of correlations in both states. In that case, the algorithmic complexity of such a procedure is *Õ* (*mn*^3^), where *Õ* bound neglects logarithmic multipliers. If we compute p-values of the tests using the permutation method, then the complexity increases to *Õ* (*mn*^3^). Thus, we use C++ language to implement the discussed analysis, significantly reducing computational time and space consumption compared to the previous methods using interpreted programming languages like Python and R. However, we developed a convenient Python wrapper of C++ functions using Numpy [23] and Pybind [24] modules. Finally, all analysis steps are efficiently parallelized, reducing computational time.

## Used datasets

### TCGA-PRAD

RNA-seq and miRNA-seq read count tables were downloaded from GDC portal (https://portal.gdc.cancer.gov/). Following the corresponding clinical description, 50 samples of the dataset were referred to “Normal” (“Solid Tissue Normal”) group and 437 samples to “Tumor” group (“Primary Tumor”). All “Tumor” tissues paired to “Normal” samples were removed to ensure the independence of samples.

Library size normalization of RNA-seq data was conducted using edgeR TMM algorithm implementation [25], and the default low-expressed gene removal procedure was applied. After normalizing for transcript length, TMM-normalized fragments per kilobase of transcript per million mapped reads (TMM-FPKM) tables were generated. Finally, TMM-FPKM tables were log_2_-transformed.

Processing of miRNA-seq data was started from the 5’-isomiR annotation from “*.mirbase21.isoforms.quantification.txt” files. For that, pri-miRNA genomic coordinates and local coordinates of canonical mature miRNA sequences were extracted from miRBase v21 [26]. The standard 5’-isomiR nomenclature was utilized: a number after “|” character denotes the shift from the canonical 5’-end in 5’-3’ direction. For example, has-miR-192-5p|+1 differs from the canonical has-miR-192-5p miRNA by the absence of the first nucleotide on its 5’-end. Note that some authors also use an additional “|” symbol and a number to simultaneously annotate 5’- and 3’-end shift [27]. This is not our case since we only considered variations at the 5’-end of miRNAs.

The following procedure was used for miRNA-seq data filtering and normalization. First, we divided each 5’-isomiR reads count by the library size of the respective sample to obtain reads per million mapped reads (RPM) tables. For a particular sample, these values reflect the proportions of reads mapped to different 5’-isomiRs. Then, low expressed isomiRs were filtered out by the default edgeR procedure. It is well known that the overwhelming majority of miRNA sequencing reads correspond to dozens of the most highly expressed isomiRs [28, 29]. Given that we marked a minimal set of 5’-isomiRs accounting for 95% of sequencing reads in a TCGA-PRAD project as highly expressed. A typical representative read distribution curve is presented in Figure 9 of “Supplementary materials”. Using this procedure, we selected 38 isomiRs from the initial pool of size 402.

Since the canonical mechanism of miRNA-mediated gene silencing is possible only in the presence of Argonaute 2 (AGO2) proteins [30, 31] we then estimated the expression levels of AGO2-loaded isomiRs by the following formula:

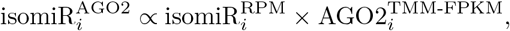

where *i* denotes the sample. We assumed that mRNA expression levels of AGO2 are proportional to the protein concentration. In the same time, miRNA-specific Argonaute loading constants did not influence the validity of our analysis since correlations between miRNAs and their targets were calculated independently for each miRNA. Previously, a similar technique significantly enhanced the predictive power of regression-based miRNA target prediction [32, 33]. As in the case of mRNA expression data, the obtained miRNA expression units were log_2_-transformed.

We used miRDB v6.0 [34] in custom prediction mode to predict targets of 5’-isomiRs (as recommended by the tool authors, interactions with target scores ≥ 80 were considered). Median expressions of the analyzed isomiRs and their predicted targets are shown in Figure 10 of “Supplementary materials”.

#### MiRNA microarray analysis of blood plasma samples

The collection of plasma specimens was formed at P.A. Herzen Moscow Oncology Research Institute. Whole blood samples was collected from 5 healthy volunteers (“Control”), 40 patients of “Low” risk group, 16 “Intermediate” risk, and 91 “High” risk group rated according EAU-ESTRO-SIOG Guidelines on Prostate Cancer [35] (see Table 2 of “Supplementary materials”). The plasma was separated according to a previously designed protocol, minimizing hemolysis and miRNA release from blood cells [36]. The level of hemolysis was evaluated by spectrophotometry [37].

Total RNA was isolated from 200 *μl* plasma by guanidine-thiocyanate-phenol-chloroform extraction with subsequent adsorption on silicon membranes using miRNeasy Serum/Plasma Kit (Qiagen) according to the instruction.

RNA was analyzed using GeneChip miRNA 4.0 (Affymetrix), detecting all noncoding and minor RNA from miRBase database [26], including mature miRNA and pre-miRNA. CEL files were processed using the oligo R package [38] in the default mode to obtain the normalized expression values.

#### Synthetic dataset to test aggregation scores

It is widely known that sample size may affect the power of statistical inferences. We study how *p*-values of the aggregation scores based on *Z_R_*(*N, T*) statistics depend on sample size and correlations in two groups. To assess the dependence, we generate two random samples of shape (*m, n*) from a multivariate normal distribution with zero means and given covariance matrices

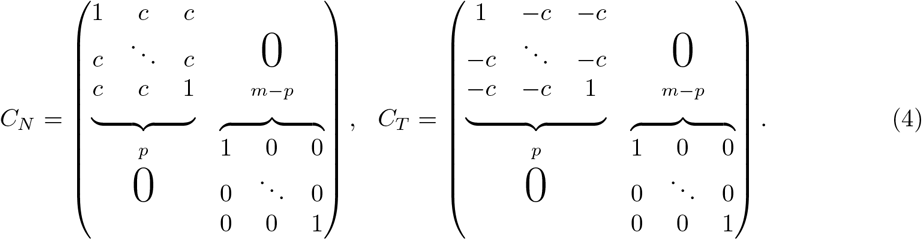

This synthetic dataset models expressions of *m* genes collected from two groups of size *n*. Using matrix *C_N_* we simulate a gene interaction network where expression of the first gene positively correlates with expressions of the following *p* — 1 genes. As seen, correlations between the first gene and other *p* – 1 genes take an opposite value in the network based on *C_T_*.

During the experiment, we fix a level of correlation *c*, percentage of changed correlations *p*, and change sample size *n* computing the mean aggregation score of the first gene. We repeat each experiment for 10 times for a fixed sample size *n*.

#### Synthetic dataset to test permutation p-value method

Comparing DCoNA with DGCA [10], we found out that the p-value computation of *Z_R_* (*N, T*) statistic implemented in DGCA is incorrect in some situations. While DCoNA recomputes *Z_R_*(*N,T*) scores after relabeling and compares the new values with the initial ones separately by genes, DGCA compares each initial score with all recomputed values. To show that the last procedure is incorrect, we generate a synthetic dataset similar to the previous one. However, we take an identity matrix of size 2m as *C_N_* and a blockdiagonal matrix as *C_T_*. More precisely, *C_T_* has p blocks of the form 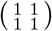 and *m* — *p* blocks of the form 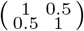. The first *p* blocks correspond to *p* gene pairs, which correlation changed more than the correlation of the remaining *m* — *p* ones. Outside these pairs, gene expressions do not correlate.

During the experiment comparing two permutation p-value computation techniques, we fix the size of “Normal” group (100 elements) and iterate the size of “Tumor” group from 10 to 100. For each covariance matrix *C_N_*, we generate the synthetic normal dataset and compute the p-value of the last pair of genes with covariance matrix of the form 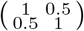 using DCoNA and DGCA procedures. We repeat each experiment for 20 times.

## Results

### DCoNA tool

We developed an easy-to-use command-line tool that performs differential co-expression network analysis on omics data. Given the expression dataset under two conditions (for example, “Normal” and “Tumor”), DCoNA identifies significantly changed correlations among all predefiend interaction pairs (network) of the form (source molecule, target molecule).

As an input DCoNA tool expects the following data (see Figure 8 of “Supplementary materials”): a table containing an expression of mRNAs, miRNAs, isomiRs, or other molecules taken under two conditions (rows of the table are indexed by the molecules while patients index columns); a table describing which of the two groups each patient belongs to; a table of the interaction pairs whose changes in correlation DCoNA will track (if the table is not given, then the described analysis applies to all possible pairs).

Aside from the pairwise correlation comparison, DCoNA can compute the aggregation scores of source molecules. These scores are designed to distinguish the source molecules with a large proportion of significantly changed correlations.

Among implementation features of DCoNA we emphpasize that it is an open-source project (https://github.com/zhiyanov/DCoNA) whose core modules are implemented in C++ language, and thus DCoNA is extremely fast. Moreover, DCoNA can be used in parallel, further speeding up the analysis. Finally, DCoNA has a user-friendly interface since it was developed as a command-line tool and Python library.

### Synthetic data analysis

#### Aggregation scores

As we described previously, we implemented several aggregation scores to track correlation changes associated with a particular gene.

The first score is based on hypergeometric test. It checks the number of significantly changed interactions with the gene to be overrepresented among other significant changes.

Other scores are based on mean, median, and other statistics of *Z_R_*(*N, T*) values associated with the gene. Using them, we compute permutation *p*-values to detect genes with large amounts of significantly changed correlations. We studied efficiency of such a procedure building the synthetic dataset (see “Materials and methods”). This dataset simulates expressions of *m* = 1000 genes: *p* of them significantly alter correlation with the first gene, while other *m* — *p* ones do not. Increasing *p*, we enlarge the proportion of changed correlations, while the value of the correlation parameter (*c*) regulates the level of the alterations.

We studied how the sample size affects the mean aggregation score using this dataset. We expected that a simultaneous increase in sample size (*n*) of “Normal” and “Tumor” groups would increase values of *Z_R_*(*N,T*) statistics and the resulting aggregation score (mean of the statistics) would consistently detect the modifications.

Figure 3 shows that the mean aggregation score of a particular gene is significant (*p*-value < 0.05) not only for large sample sizes but also for small ones depending on the level of correlation (*c*). For example, if a significant part (10%) of interactions changes, the mean aggregation score will detect these alterations even for 20 elements in both groups.

**Figure 3:**
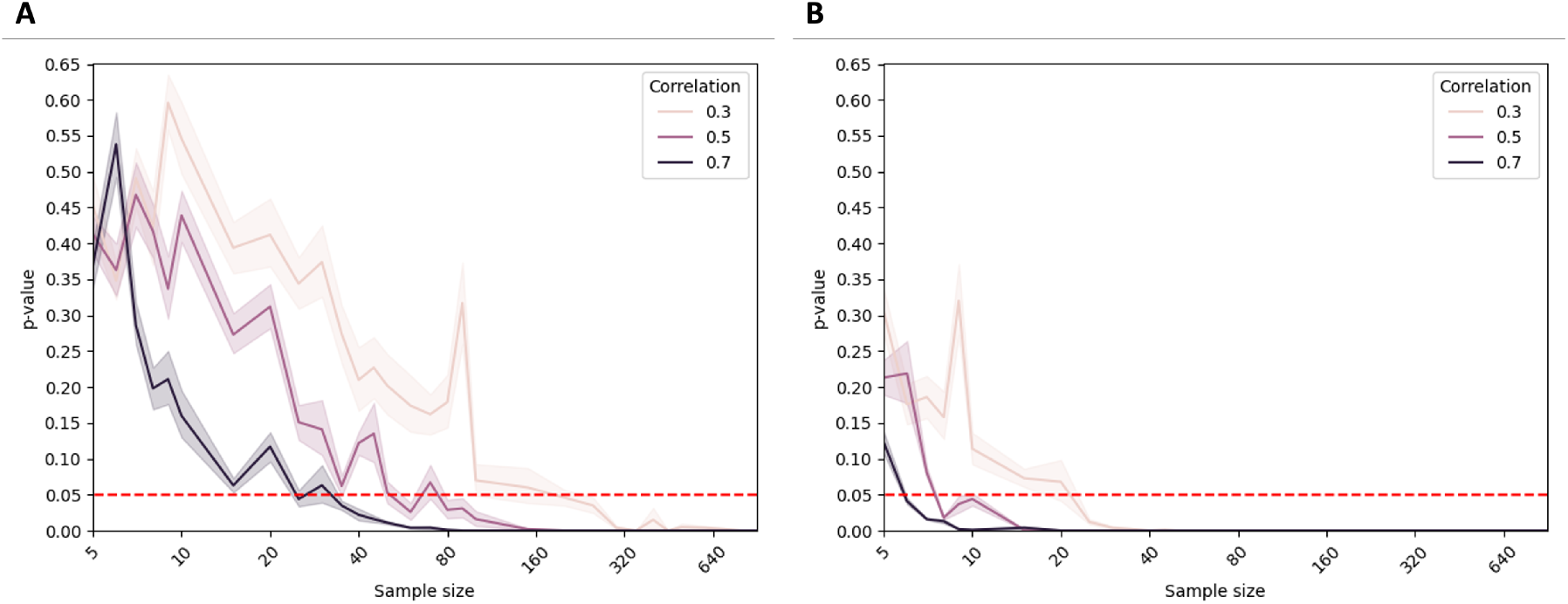
Dependence of *p*-value of the mean aggregation score on the sample size. Only *p* percent of *m* = 1000 genes change its correlation with another gene from the value *c* to —*c*. The experiment was repeated 10 times to evaluate the standard deviation.

#### Comparison with DGCA: permutation *p*-value

We compared DCoNA and DGCA *Z_R_*(*N,T*) permutation *p*-value computation approaches. The common part of both methods is that they shuffle labels of the groups and recompute *Z_R_*(*N,T*) statistics. However, unlike DGCA, DCoNA compares the new values with the initial ones separately by genes.

To assess both procedures, we used a synthetic dataset similar to the previous one (see “Materials and methods”). This dataset simulates expression of 20 genes using multivariate normal distribution. Genes of this dataset break down into *m* = 10 independent pairs.

During the experiment, correlations of *p* gene pairs change more (from 0 to 1) than the remaining correlations (from 0 to 0.5). For a small sample size of “Tumor” group (in comparison to “Normal” one), even after label shuffling, we expected *Z_R_*(*N,T*) to be larger for those pairs of genes, which correlations changed more. This effect would allow us to distinguish DGCA and DCoNA procedures of permutation *p*-value computation on a pair with a relatively small change in correlation.

As Figure 4 shows, DCoNA *p*-value computation approach is more accurate than one proposed in DGCA tool, especially when a large proportion (*p* = 0.9) of correlations changed more than the rest. More precisely, for the sample size values of “Tumor” group in (40, 70) interval, the hypothesis on correlation equivalence is accepted by DGCA and rejected by DCoNA tool.

**Figure 4:**
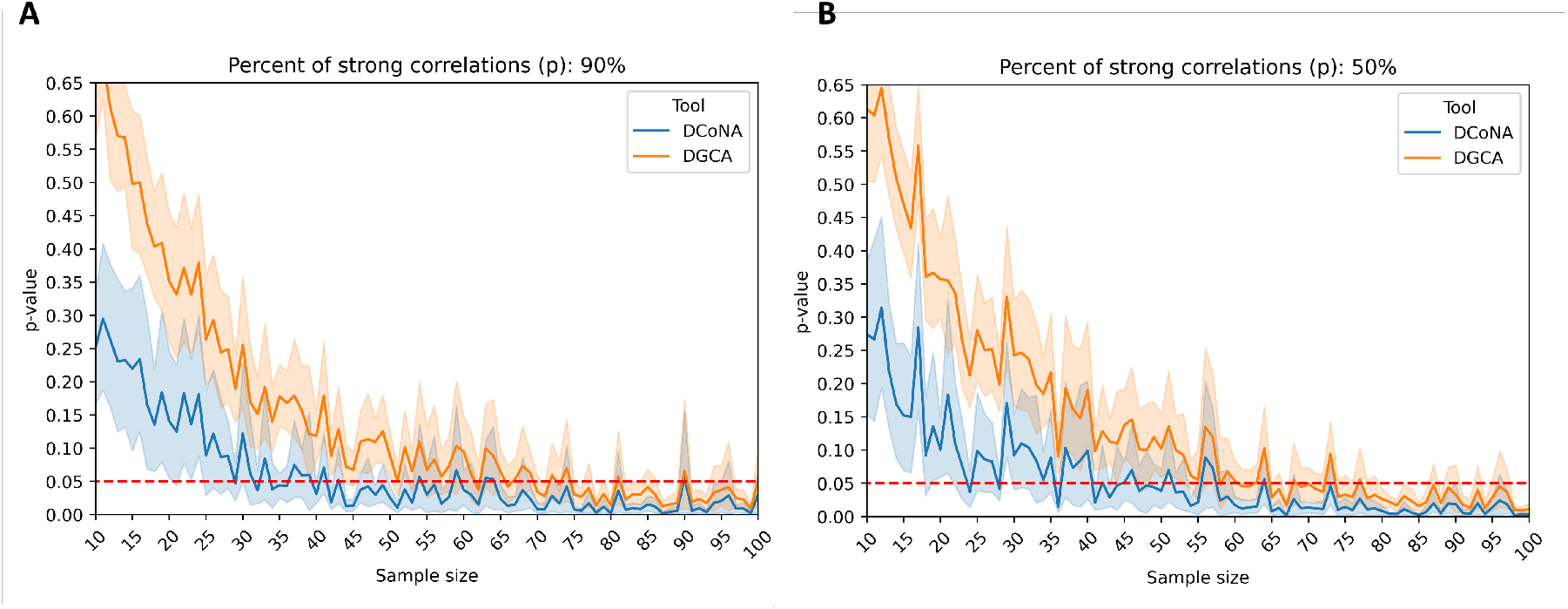
Comparison of the permutation *p*-value computation approaches of *Z_R_*(*N, T*) test between DGCA and DCoNA. The sample size of “Normal” group is equal to 100, while the size of “Tumor” group changes. The experiment was repeated 20 times to estimate the standard deviation.

### Prostate cancer data analysis

#### Comparison with DGCA: time and space consumption

To compare DCoNA and DGCA in terms of time and memory cosumption, we used TCGA-PRAD dataset.

In the first experiment, we changed the number of tested genes and analyzed all pairs composed of these genes to be differentially correlated. We launched both tools to compare Spearman correlations with the number of genes from 100 to 3000 with a step of 50. Theoretically, with the number of genes *n*, both tools perform *O*(*n*^2^) operations during the analysis. However, in Figure 5 A, we see that DCoNA is at least 100 times faster than DGCA (even in one thread mode). Similar trend is true for RAM usage (Figure 5 C).

**Figure 5:**
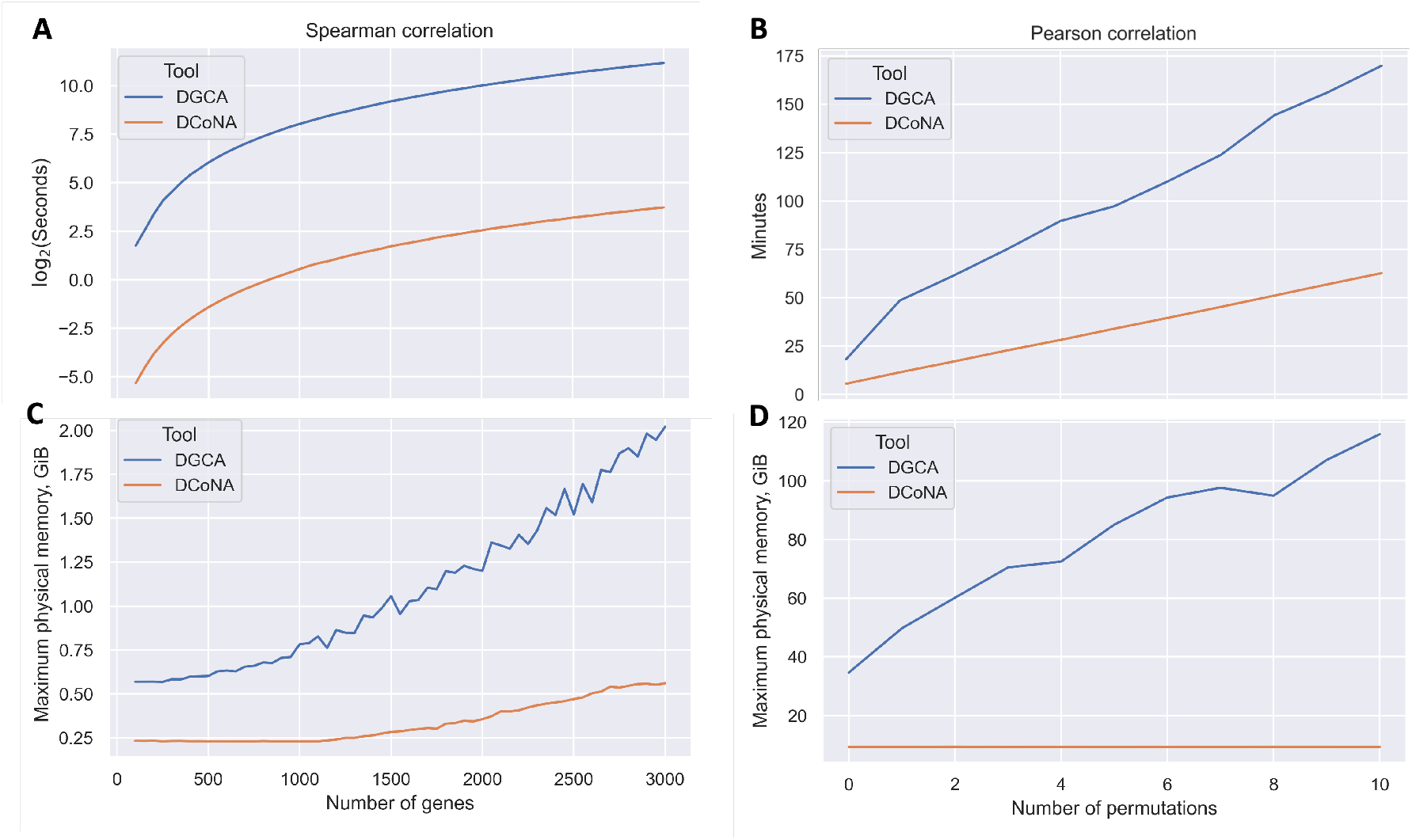
Comparison of time and space consumption of DCoNA and DGCA. **A, C**: dependence of resource consumption on the number of analyzed genes; **B, D**: on the number of permutations in the permutation *p*-value computation procedure.

In the second experiment (see Figure 5 B, D), we fixed the number of tested genes (15000) and performed the previous analysis with one difference. Instead of the analytical *p*-value computation procedure, we used the permutation one. During DGCA execution RAM usage exceeded 100GiB threshold. In contrast, in the case of DCoNA, RAM usage did not increase (< 8GiB) with the number of permutations, so it can be executed even on personal computers.

#### Targetome of 5’-isomiRs in healthy prostate

Using DCoNA we aimed to study how co-expression of pairs of types (isomiR, mRNA) changed in prostate cancer. In the first step of the analysis, we extracted 38 highly expressed isomiRs, accounting for 95% of the total isomiR expression (Figure 9 of “Supplementary materials”). As seen from the figure, there are 5 highly expressed non-canonical isomiRs (their seed regions differ from the canonical ones).

Next, we built a bioinformatically predicted regulatory network of the highly expressed isomiRs and their target mRNAs using miRDB tool [34]. Further, we selected the interactions with Spearman correlation of expression levels less than −0.3. The same standard criterion was previously used in many works [39, 40, 41]. The resulting predicted regulatory networks are shown in Figure 6: as we have already discussed, isomiRs of one miRNA correlate with different targets.

**Figure 6:**
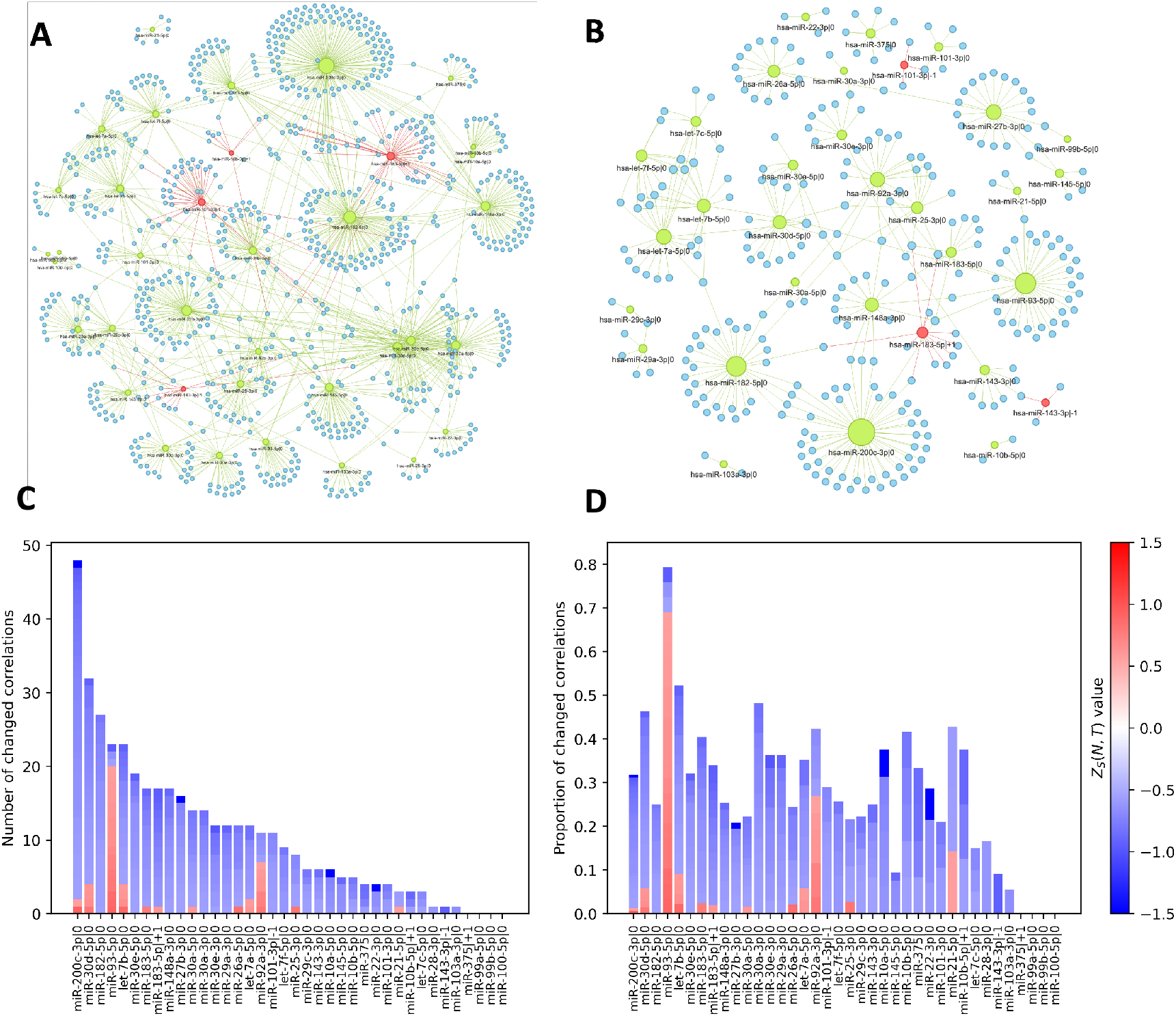
Predicted regulation networks of isomiRs and their target mRNAs in **A**: “Normal” state; **B**: “Tumor” state of TCGA-PRAD dataset. **C**, **D** represent the number of significantly changed correlations that jumped over —0.3 threshold: blue bars represent the loss of negative correlation while red ones represent new negative correlations. **C**: the absolute number of changed correlations, **D**: number of changed correlations normalized by the number of targets in “Normal” state.

#### Systematic disregulation of 5’-isomiR targeting in prostate cancer

Among other things, Figure 6 shows that a large part of the highly expressed isomiRs lost their targets in “Tumor” state. To study this effect deeper, we performed the following analysis. Using DCoNA, we tracked those target mRNAs wich Spearman correlation with source isomiRs statistically significantly “jumped” over —0.3 threshold. Figure 6 C, D reveals a general and powerful trend: for almost all highly expressed isomiRs the number of new targets in “Tumor” state is less than the number of lost ones. The exceptions are hsa-miR-93-5p|0 and hsa-miR-92a-3p|0 molecules. In “Normal” state they had 29, 26 gene-targets (with correlation < —0.3), while in “Tumor” state 3, 4 of them disappeared and 20, 7 appeared, respectively.

Further, we applied DESeq2 [7] to our data to check whether the regulation loss is connected with the change of molecules expression. Interestingly, expression of hsa-miR-93-5p|0 and hsa-miR-92a-3p|0 significantly increased from normal tissues to the cancer ones: log_2_(fold change) = 1.77 (adjusted *p*-value = 8.7*e*–66) and log_2_(fold change) = 1.09 (adjusted *p*-value = 6.7*e*–34), respectively. However, the other isomiRs with increased expressions (14 molecules of 16) did not have a significant increase in targets. For example, 6 (including the most increased one) of them had no new targets at all. Simultaneously, 16 of 17 isomiRs with significantly decreased expression did not acquire new targets too. For more details see Table 1 of “Supplementary materials”.

#### High expression of hsa-miR-93-5p is associated with cancer aggressivness

Among the newly appeared targets of hsa-miR-93-5p|0, we highlight FBXO31 and MFN2. Concordantly, their expression significantly reduced in prostate cancer: log_2_(fold change) = —0.48 and — 0.66, respectively. In addition, these molecules were not negatively correlated with any other predicted isomiR regulators in “Tumor” state. Thus, these targets may play a tumor suppressive role in prostate cancer (see Discussion for details).

Also, we checked hsa-miR-93-5p|0 targets to be experimentally supported using DIANA-TarBase database [42]. It turned out that 20 of 29 and 21 of 35 bioinformatically predicted mRNA targets are experimentally validated in “Normal” and “Tumor” states, respectively.

Given the predicted oncomiR role of hsa-miR-93-5p|0 (elevated expression and appearance of cancer-specific targets), we analyzed whether the expression of hsa-miR-93-5p|0 varied between tumors with different aggressiveness. Using a clinical description of TCGA-PRAD samples, we divided “Tumor” samples into three groups depending on Gleason’s score: “Low” risk group with Gleason’s score ≤ 6 (40 samples), “Intermediate” risk group with Gleason’s score = 7 (203 samples), and “High” risk group with Gleason’s score ? 6 (194 samples). As a “Control”, we took samples from “Normal” group (50 samples). As seen in Figure 7, the isomiR’s expression increased from “Control” to “Low” and from “Low” to “High” risk groups (in both cases Mann-Whitney *U*-test *p*-value < 0.01).

**Figure 7:**
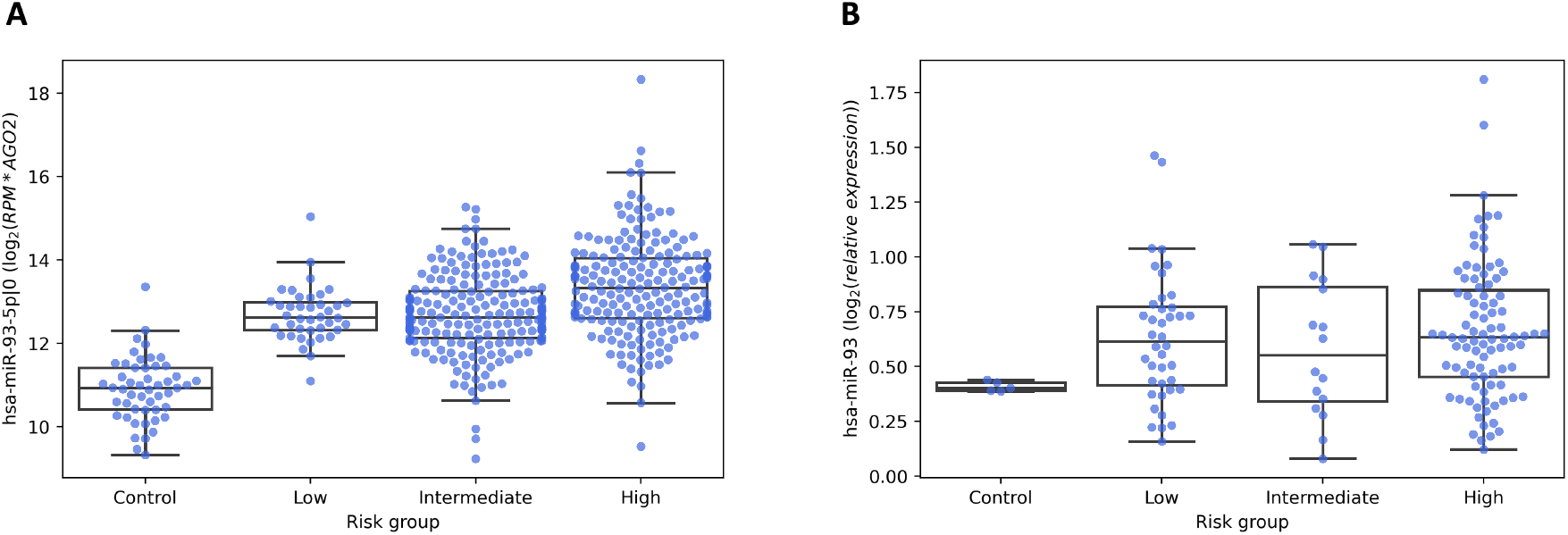
Expression of hsa-miR-93 grouped by Risk Groups (Gleason’s score) in **A**: blood plasma and **B**: TCGA-PRAD. In both cases, the expression in “Healthy” group significantly differed from “High” risk group.

The aforementioned analysis was conducted with TCGA primary tumor samples. To extend our findings beyond primary tumors, we analyzed miRNA expression profiles in blood plasma samples derived from 147 patients with prostate cancer and 5 patients without any prostate pathologies (for more details, see Table 2 of “Supplementary materials”). In agreement with TCGA data, hsa-miR-93 expression significantly differed between “Control” and “High” risk groups (Mann-Whitney *U*-test *p*-value < 0.05). At the same time expression of other miRNAs in our analysis (Figure 9 of “Supplementary materials”) did not change significantly between these groups (*p*-value > 0.05). This fact additionally emphasizes hsa-miR-93 among other miRNAs as a potential marker of aggressive prostate cancer.

## Discussion

We developed DCoNA - the user-friendly command-line tool and Python library identifying differential co-expression interactions in omics data. Given two states (for example, “Normal” and “Tumor”), DCoNA checks all predefined interactions to have different correlations in these states. Note that the primary analysis comparing correlations may lack rigor: random fluctuations in expression can change correlation values. Hence, DCoNA uses a particular statistical test based on Fisher *z*-statistic to test a hypothesis on correlation equivalence. Aside from the differential correlation analysis, DCoNA can be used to distinguish those regulatory nodes whose interactions significantly changed overall. Such nodes may play a significant role in distinguishing the states and be the subject of separate studies.

To the best of our knowledge, the current state-of-the-art algorithm providing similar analysis is DGCA [10]. We compared DCoNA with the analog using several synthetic examples and TCGA-PRAD (prostate cancer) dataset. We proved that DCoNA is significantly faster and can be used not only on computational servers but also on personal computers: DCoNA consumes considerably less RAM than DGCA. Also, in contrast to DCoNA, DGCA can not be applied to a predefined regulatory network and performs its analysis for all molecules’ pairs of a dataset, significantly reducing the number of applications (one of them will be considered further). Finally, the procedure of the permutation *p*-value computation implemented in DGCA is incorrect in some cases, which has been corrected in DCoNA.

We applied DCoNA to the regulatory network of miRNA isoforms (isomiRs) and their bioinformatically predicted mRNA targets using TCGA-PRAD dataset. It turned out that the most highly expressed isomiRs lost their negative correlation in “Tumor” samples compared to “Normal” ones. hsa-miR-93-5p|0 was an exception: the number of the new targets with negative correlation in “Tumor” state was essentially greater than in “Normal” state. About two-thirds of these targets turned out to be experimentally confirmed. Among the new targets of the isomiR, we highlight FBXO31 and MFN2. FBXO31, the member of protein-ubiquitin ligase, activates ERK- and su-presses PI3K-AKT-mediated signaling pathways in prostate cancer by promoting the degradation of DUSP6 [43]. The direct interaction between miR-93 and 3’-UTR of FBXO31 was experimentally validated in several breast cancer cell lines by Manne et al. [44]. Another prostate cancer-specific target of miR-93, MFN2, was shown to suppress breast and thyroid cancer progression through AKT signaling [45, 46]. The direct binding of miR-93 to 3’-UTR of MFN2 was also experimentally validated with luciferase reporter assays [47, 48].

We also found that expression of hsa-miR-93-5p|0 was associated with the cancer aggressiveness: TCGA primary tumor samples with higher Gleason’s score had higher expression of the isomiR. We confirmed this by analyzing blood plasma samples (153 patients) using Affymetrix miRNA microarrays. Consensus with TCGA-PRAD, expression of miR-93 increased in “High” risk group compared to “Control” group: other highly expressed isomiRs did not show this trend. This fact also highlighted hsa-miR-93-5p|0 as isomiR playing a clear oncogenic role in prostate cancer.

Similar results in deregulation of isomiR-target co-expression networks were recently obtained by Telonis et al. [49]. Specifically, the authors reported a large-scale “differential wiring” of isomiRs and putative targets between normal breast tissues and triple-negative breast cancer. Moreover, the obtained results differed in each race. One of the possible mechanisms underlying isomiR-target differential co-expression was based on alternative splicing. For example, hsa-miR-200b-3p had the splice variant-specific binding site in one of RTN4 exons, and this RTN4 splice variant became downregulated in breast cancer tissues. Thus, mRNA isoform switching can explain differences in isomiR-target co-expression profiles. Note that such situations were not covered in our analysis, since miRDB predicted isomiR targets only within mRNA 3’-UTR sequences.

## Data Availability Statement

The data discussed in this publication have been deposited in NCBI’s Gene Expression Omnibus [50] and are accessible through GEO Series accession number GSE206793 (https://www.ncbi.nlm.nih.gov/geo/query/acc.cgi)

## Acknowledgment

The authors thank Prof. Alexey Galatenko from Faculty of Biology and Biotechnology, HSE University and Prof. Isidore Rigoutsos from Computational Medicine Center, Thomas Jefferson University, for useful comments and discussion.

## Supplementary materials

**Tables**

**Table 1:**
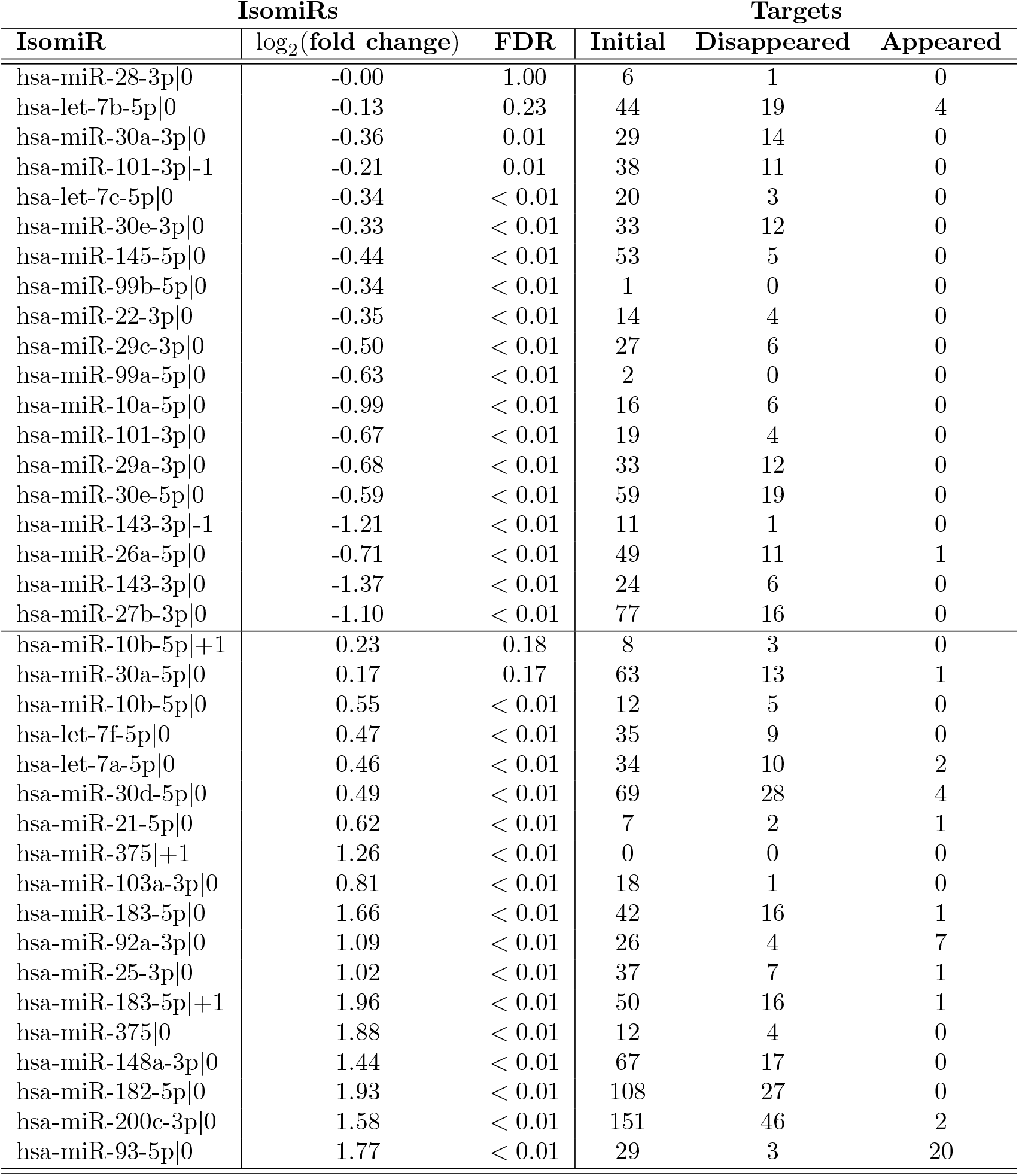
**IsomiRs**: ifferential expression of the highly expressed isomiRs in TCGA-PRAD (“Normal” vs “Tumor”). **Initial**: the number of bioinformatically predicted mRNA targets in “Normal” state with correlation < −0.3. **Disappeared**: the number of targets significantly “jumped” over the correlation threshold and lost negative correlation. **Appeared**: the number of targets significantly “jumped” over the correlation threshold and acquired negative correlation.

**Table 2:**
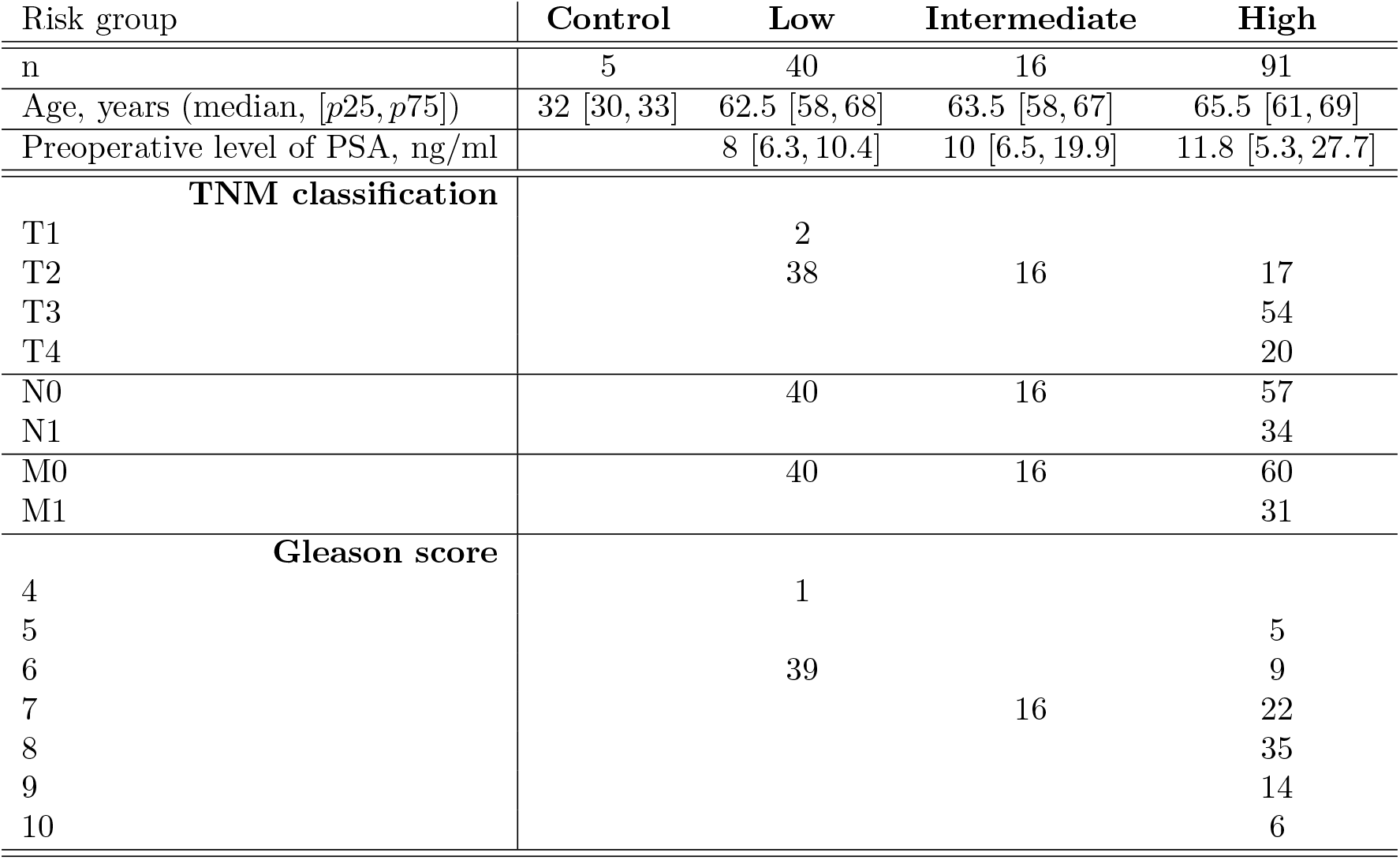
Clinical characteristics of patients included in the study.

**Figures**

**Figure 8:**
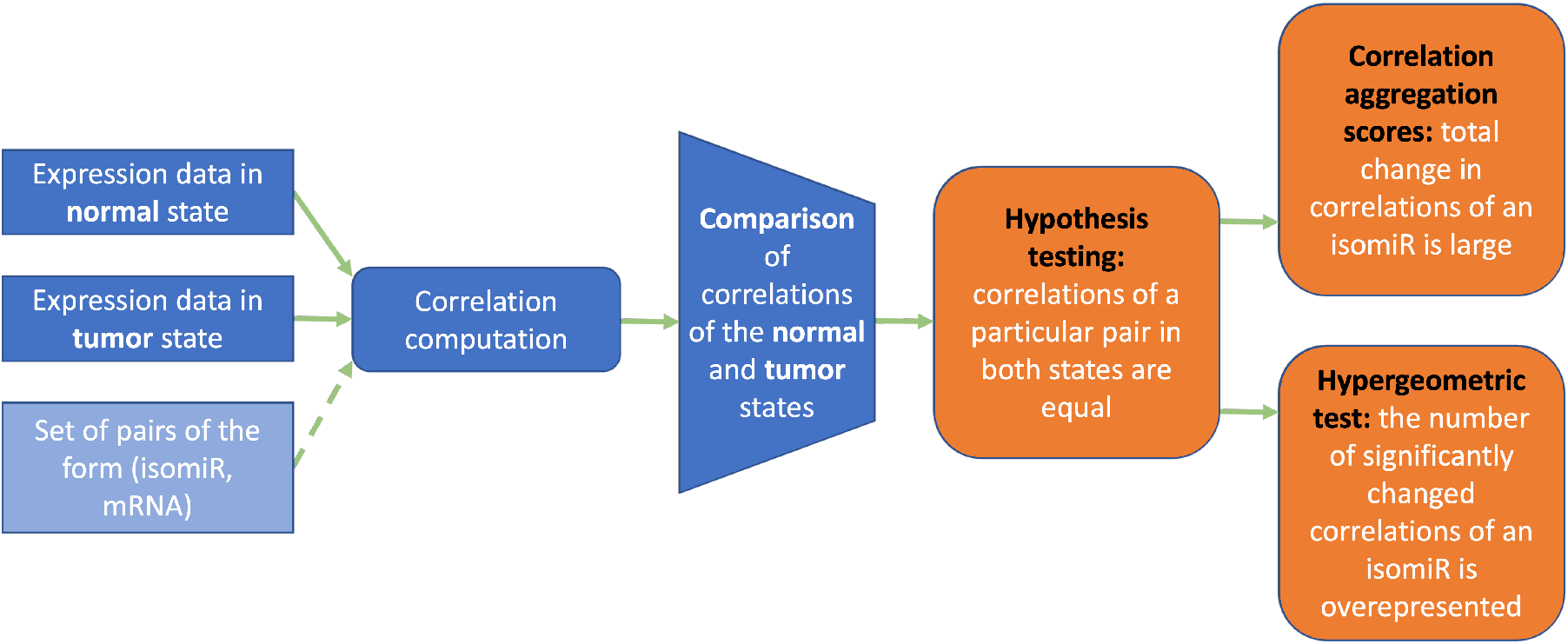
Computational pipeline of DCoNA. The dashed line represents an optional network mode. If the set of pairs of interest is not defined, DCoNA performs the differential co-expression analysis for all possible pairs of molecules from the expression data.

**Figure 9:**
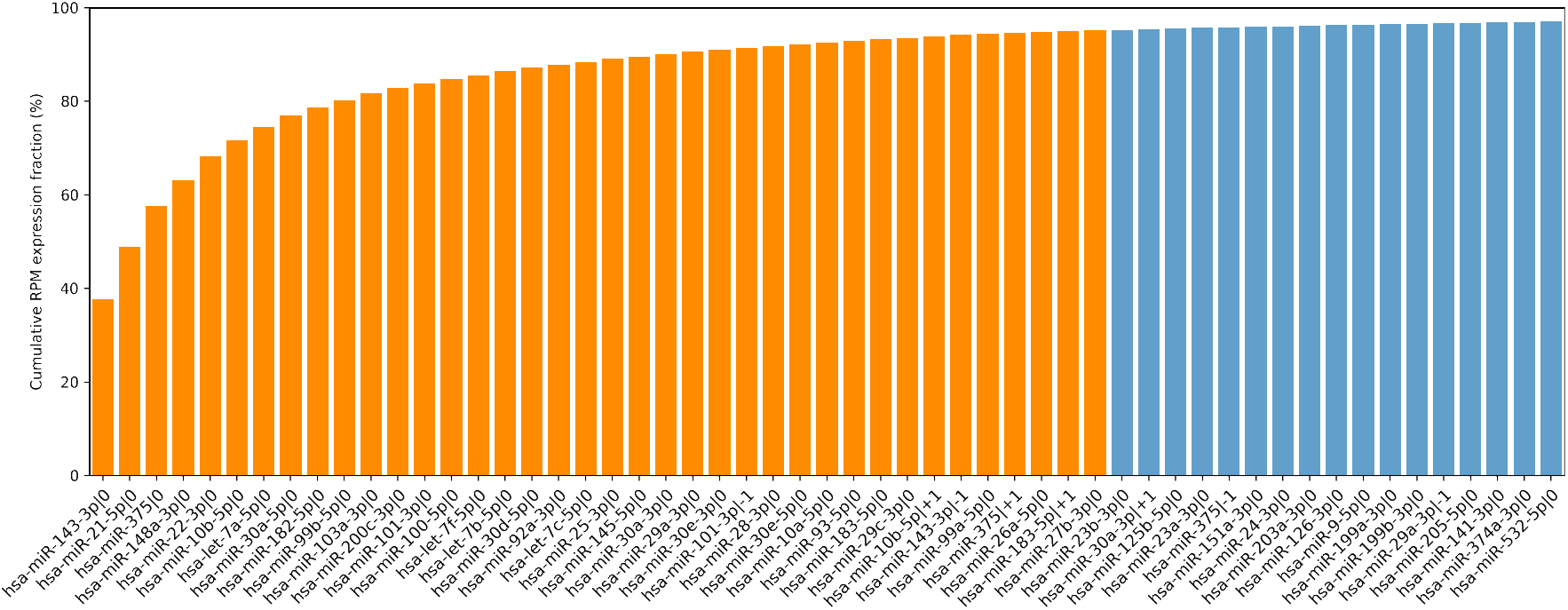
Cumulative expression fraction of isomiRs in TCGA-PRAD dataset. The orange bars define 95% threshold, i.e. 95% of expression comes from 38 isomiRs. IsomiRs are sorted in descending order of expression.

**Figure 10:**
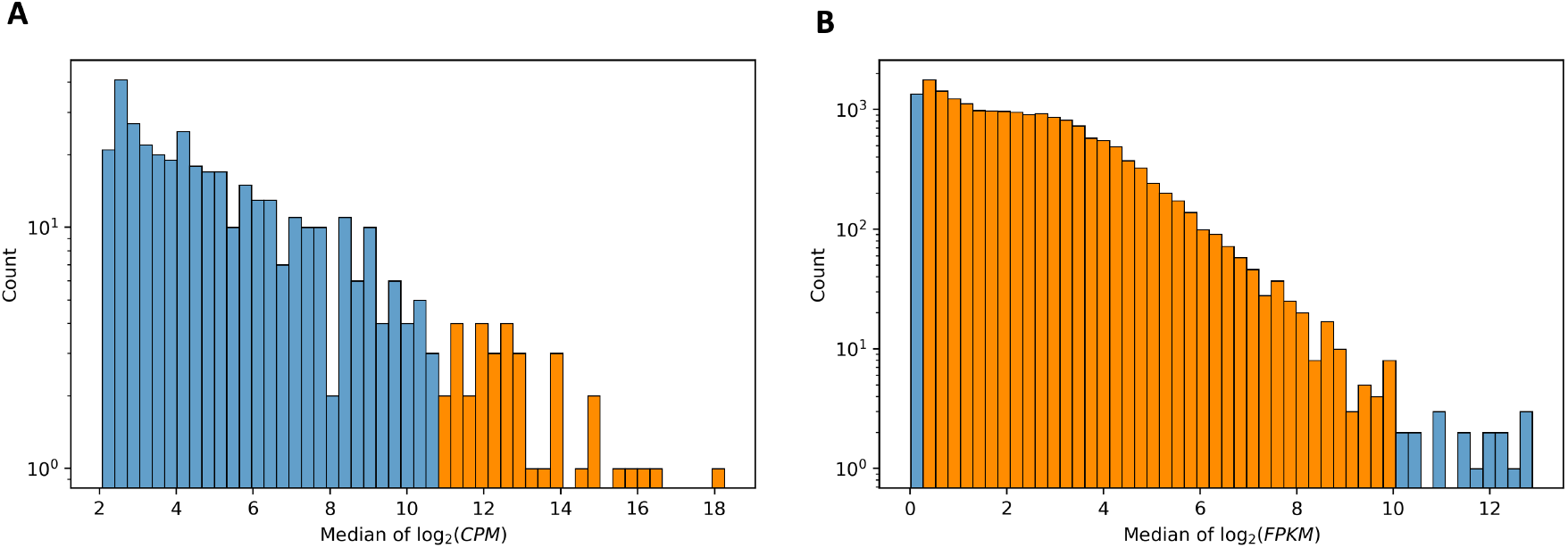
Histograms of **A**: isomiRs and **B**: mRNAs expression in TCGA-PRAD. The orange bars higlight the analised molecules: the higly expressed isomiRs and their target mRNAs (bionfirmatically predicted and validated with correlation).

