## Supplementary tables and figures for "Differential co-expression network analysis with DCoNA reveals isomiR targeting aberrations in prostate cancer"

### Supplementary materials

| IsomiRs | | | Targets | | |
|---|---|---|---|---|---|
| **IsomiR** | $\log_2$(**fold change**) | **FDR** | **Initial** | **Disappeared** | **Appeared** |
| hsa-miR-28-3p\|0 | -0.00 | 1.00 | 6 | 1 | 0 |
| hsa-let-7b-5p\|0 | -0.13 | 0.23 | 44 | 19 | 4 |
| hsa-miR-30a-3p\|0 | -0.36 | 0.01 | 29 | 14 | 0 |
| hsa-miR-101-3p\|-1 | -0.21 | 0.01 | 38 | 11 | 0 |
| hsa-let-7c-5p\|0 | -0.34 | < 0.01 | 20 | 3 | 0 |
| hsa-miR-30e-3p\|0 | -0.33 | < 0.01 | 33 | 12 | 0 |
| hsa-miR-145-5p\|0 | -0.44 | < 0.01 | 53 | 5 | 0 |
| hsa-miR-99b-5p\|0 | -0.34 | < 0.01 | 1 | 0 | 0 |
| hsa-miR-22-3p\|0 | -0.35 | < 0.01 | 14 | 4 | 0 |
| hsa-miR-29c-3p\|0 | -0.50 | < 0.01 | 27 | 6 | 0 |
| hsa-miR-99a-5p\|0 | -0.63 | < 0.01 | 2 | 0 | 0 |
| hsa-miR-10a-5p\|0 | -0.99 | < 0.01 | 16 | 6 | 0 |
| hsa-miR-101-3p\|0 | -0.67 | < 0.01 | 19 | 4 | 0 |
| hsa-miR-29a-3p\|0 | -0.68 | < 0.01 | 33 | 12 | 0 |
| hsa-miR-30e-5p\|0 | -0.59 | < 0.01 | 59 | 19 | 0 |
| hsa-miR-143-3p\|-1 | -1.21 | < 0.01 | 11 | 1 | 0 |
| hsa-miR-26a-5p\|0 | -0.71 | < 0.01 | 49 | 11 | 1 |
| hsa-miR-143-3p\|0 | -1.37 | < 0.01 | 24 | 6 | 0 |
| hsa-miR-27b-3p\|0 | -1.10 | < 0.01 | 77 | 16 | 0 |
| hsa-miR-10b-5p\|+1 | 0.23 | 0.18 | 8 | 3 | 0 |
| hsa-miR-30a-5p\|0 | 0.17 | 0.17 | 63 | 13 | 1 |
| hsa-miR-10b-5p\|0 | 0.55 | < 0.01 | 12 | 5 | 0 |
| hsa-let-7f-5p\|0 | 0.47 | < 0.01 | 35 | 9 | 0 |
| hsa-let-7a-5p\|0 | 0.46 | < 0.01 | 34 | 10 | 2 |
| hsa-miR-30d-5p\|0 | 0.49 | < 0.01 | 69 | 28 | 4 |
| hsa-miR-21-5p\|0 | 0.62 | < 0.01 | 7 | 2 | 1 |
| hsa-miR-375\|+1 | 1.26 | < 0.01 | 0 | 0 | 0 |
| hsa-miR-103a-3p\|0 | 0.81 | < 0.01 | 18 | 1 | 0 |
| hsa-miR-183-5p\|0 | 1.66 | < 0.01 | 42 | 16 | 1 |
| hsa-miR-92a-3p\|0 | 1.09 | < 0.01 | 26 | 4 | 7 |
| hsa-miR-25-3p\|0 | 1.02 | < 0.01 | 37 | 7 | 1 |
| hsa-miR-183-5p\|+1 | 1.96 | < 0.01 | 50 | 16 | 1 |
| hsa-miR-375\|0 | 1.88 | < 0.01 | 12 | 4 | 0 |
| hsa-miR-148a-3p\|0 | 1.44 | < 0.01 | 67 | 17 | 0 |
| hsa-miR-182-5p\|0 | 1.93 | < 0.01 | 108 | 27 | 0 |
| hsa-miR-200c-3p\|0 | 1.58 | < 0.01 | 151 | 46 | 2 |
| hsa-miR-93-5p\|0 | 1.77 | < 0.01 | 29 | 3 | 20 |

Table 1: **IsomiRs**: ifferential expression of the highly expressed isomiRs in TCGA-PRAD ("Normal" vs "Tumor"). **Initial**: the number of bioinformatically predicted mRNA targets in "Normal" state with correlation $< -0.3$. **Disappeared**: the number of targets significantly "jumped" over the correlation threshold and lost negative correlation. **Appeared**: the number of targets significantly "jumped" over the correlation threshold and acquired negative correlation.

| Risk group | **Control** | **Low** | **Intermediate** | **High** |
|---|---|---|---|---|
| n | 5 | 40 | 16 | 91 |
| Age, years (median, [$p25, p75$]) | 32 [30, 33] | 62.5 [58, 68] | 63.5 [58, 67] | 65.5 [61, 69] |
| Preoperative level of PSA, ng/ml | | 8 [6.3, 10.4] | 10 [6.5, 19.9] | 11.8 [5.3, 27.7] |
| **TNM classification** | | | | |
| T1 | | 2 | | |
| T2 | | 38 | 16 | 17 |
| T3 | | | | 54 |
| T4 | | | | 20 |
| N0 | | 40 | 16 | 57 |
| N1 | | | | 34 |
| M0 | | 40 | 16 | 60 |
| M1 | | | | 31 |
| **Gleason score** | | | | |
| 4 | | 1 | | |
| 5 | | | | 5 |
| 6 | | 39 | | 9 |
| 7 | | | 16 | 22 |
| 8 | | | | 35 |
| 9 | | | | 14 |
| 10 | | | | 6 |

Table 2: Clinical characteristics of patients included in the study.

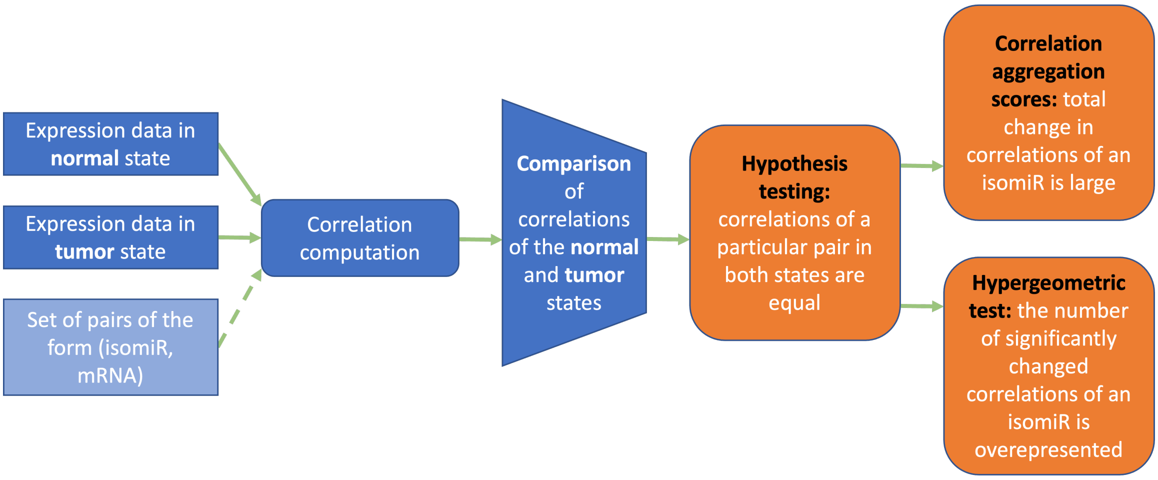


Figure 1: Computational pipeline of DCoNA. The dashed line represents an optional network mode. If the set of pairs of interest is not defined, DCoNA performs the differential co-expression analysis for all possible pairs of molecules from the expression data.

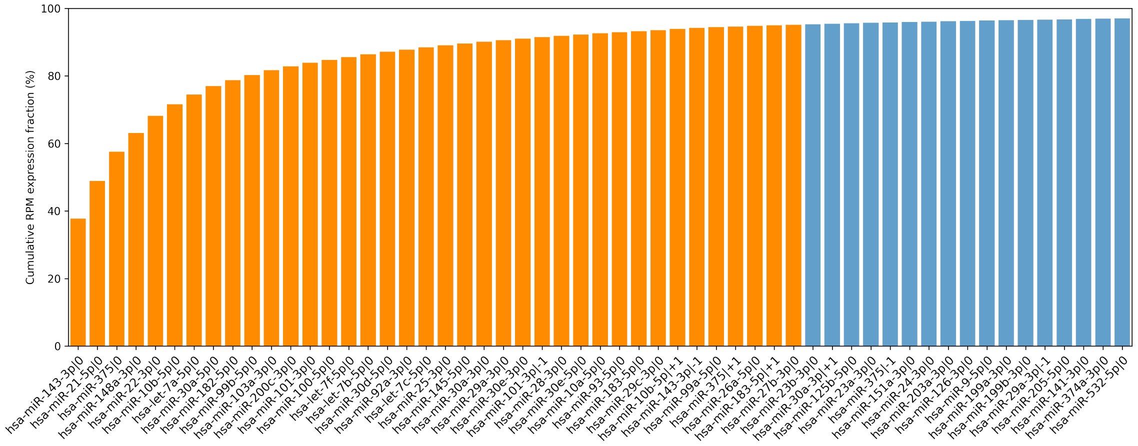


Figure 2: Cumulative expression fraction of isomiRs in TCGA-PRAD dataset. The orange bars define 95% threshold, i.e. 95% of expression comes from 38 isomiRs. IsomiRs are sorted in descending order of expression.

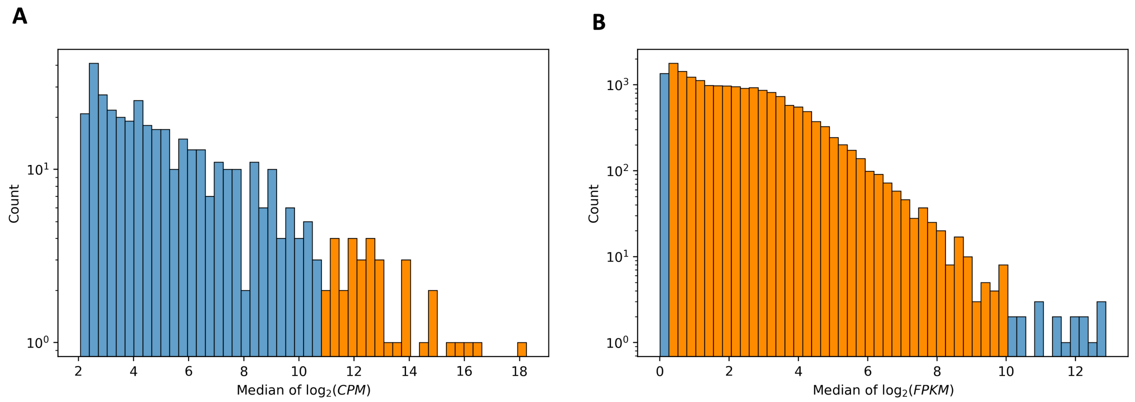


Figure 3: Histograms of **A**: isomiRs and **B**: mRNAs expression in TCGA-PRAD. The orange bars higlight the analised molecules: the higly expressed isomiRs and their target mRNAs (bionfirmatically predicted and validated with correlation).
